# Or22 allelic variation alone does not explain differences in discrimination of yeast-produced volatiles by *D. melanogaster*

**DOI:** 10.1101/186064

**Authors:** Carolyn Elya, Allison S. Quan, Kelly M. Schiabor, Michael B. Eisen

## Abstract

Different lines of the fruit fly *Drosophila melanogaster* show variation in the ability to discriminate between volatiles produced by the yeast *Saccharomyces cerevisiae* under natural (nitrogen-limiting, YVN) or laboratory (sugar-limiting, YVL) conditions. Previous work in our laboratory uncovered a strong correlation between heightened sensitivity to YVN wild *D. melanogaster* lines that harbored a chimeric variant of the highly variable odorant receptor 22 (Or22) locus of *D. melanogaster*. We sought to determine if this trend held for an extended set of *D. melanogaster* lines, if observed variation within chimeric and non-chimeric lines could be explained by nucleotide polymorphisms and if replacing Or22 with a chimeric allele in a non-chimeric background could confer the enhanced ability to detect YVN. In parallel, we performed crosses of chimeric and non-chimeric fly lines and assayed the behavior of their progeny for enhanced sensitivity to YVN to assess the heritability of the Or22 locus. Ultimately, we found that, while the overall trend of increased sensitivity to YVN in chimeric lines persists, there are exceptions and variation that cannot be explained by sequence variation at the Or22 locus. In addition, we did not observe increased sensitivity for YVN upon replacing the Or22 allele in a non-chimeric line (OreR) with that from our most YVN-sensitive, chimeric line (ME). Though our results do not support our hypothesis that Or22 is the primary driver of sensitivity to YVN, Or22 remains an interesting locus in the context of fly-yeast ecology.

## Introduction

The fruit fly *Drosophila melanogaster* and the yeast *Saccharomyces cerevisiae* are close natural partners: flies require yeast for development and nutrition (1) and yeast depend on flies to be vectored to new substrates (2). Flies can sense a variety of compounds that are produced by fermenting yeast via olfaction (3) and have demonstrated a preference to yeasted over non-yeasted fruit in the context of the laboratory (4). Evidence to date suggests that chemical communication is the basis of the co-occurrence of flies and yeast in nature, but the specific components that mediate this molecular conversation are incompletely understood.

Olfactory sensing in *Drosophila* begins in the antenna and maxillary palp, the two main odor sensing organs in adult flies (5). Both the antenna and maxillary palp are covered with sensory hairs (sensilla) which house one to four olfactory receptor neurons (6). These neurons express transmembrane odorant receptors and project onto distinct glomeruli in the antennal lobe, the central olfactory processing center (6). Olfaction is sensed when a volatile compound (odorant) diffuses into a sensillum and binds its cognate olfactory receptor (3) thereby eliciting a stimulus that is processed by the antennal lobe (6).

The *Drosophila* genome encodes 62 different olfactory receptors, each of which is expressed in a particular type of olfactory neuron either alone or in conjunction with up to two additional olfactory receptor types (3, 7). All neurons expressing a given olfactory receptor project onto the same glomerulus within the antennal lobe (6). Extensive work has profiled the repertoire of each odorant receptor by recording responses of neurons ectopically-expressing olfactory receptors to a panel of 110 odors, revealing that *D. melanogaster* odorant receptors can detect a diverse set of organic compounds with varying sensitivity and response kinetics (8).

Previous work in our laboratory showed that the wild-type fly line Ral437 (9) can differentiate between volatiles produced by yeast under natural (nitrogen-limiting, YVN) or laboratory (sugar-limiting, YVL) conditions and that six volatile compounds mediate this attraction (10). Three of these compounds, ethyl hexanoate, ethyl octanoate, and isoamyl acetate, are recognized by the same odorant receptor, Or22a (8, 11). Intriguingly, genomic comparison of *Scaptomyza flava*, an herbivorous drosophilid, and *D. melanogaster* found that Or22a is one of two olfactory receptors conserved among drosophilids but completely lost in *S. flava*, suggesting that Or22a plays a role in the fungivorous lifestyle of *D. melanogaster* (12).

In *D. melanogaster*, Or22a is one of two tandem copies of Or22 (the other copy being Or22b) present at the Or22 locus on chromosome 2L (13). Both odorant receptors are expressed in basiconic sensilla of the ab3A olfactory neuron (3, 14). A tandem duplication of Or22 occurred in the *D. melanogaster* lineage prior to the divergence from *D. simulans* but after divergence from the *D. erecta* and *D. yakuba* lineage (13). The Or22 locus is functionally variable between *Drosophila* species, indicating that it is a quickly evolving region and likely under selective pressure (15, 16). In *D. erecta*, Or22 has evolved to sense odors from the host plant *Pandanus* spp (17). In *D. sechellia*, Or22a has specialized to detect odors that emanate from the host plant *Morinda citrifolia* while Or22b has decayed into a pseudogene (18).

In addition to being highly variable between species, studies have observed significant sequence variability at the Or22 locus between different lines of *D. melanogaster* (13, 19). A set of *D. melanogaster* lines were found to segregate by two variants at the Or22 locus: one non-chimeric variant contained two copies of Or22, Or22a and Or22b, while the other contained a chimera (Or22ab) consisting of the first exon of Or22a fused to the last three exons of Or22b (13). In addition to the length variants observed in *D. melanogaster* lines, some lines were also observed to possess an inversion on 2L whose breakpoint is just 0.7 Mb away from the Or22 locus; however, no association between the inversion and the length variant was observed (13). Despite tolerating substantial variation, the Or22 locus has been implicated as a region undergoing positive selection in comparative population genetic studies of *D. melanogaster* in African and Europe (20). In Australia, the presence of the length variants is clinal, where all southern lines were non-chimeric at the Or22 locus and almost all northern flies were chimeric (19). These studies suggest that the Or22 locus has recently undergone positive selection with *D. melanogaster*, though the conferred benefit indicated by that selection is unknown (13).

Previous work in our laboratory had demonstrated a strong correlation between sensitivity to YVN over YVL and the chimeric allele of Or22 using a trap-based olfactory assay (21). We hypothesized that the chimeric variant of the Or22 locus confers a heightened sensitivity to differences in yeast volatile bouquets and consequently contributes to flies’ ability to locate yeast in nature. Here, we sought to explore this hypothesis by expanding our behavioral set with wild, inbred lines, assaying the behavior of progeny of reciprocal crosses from this set, analyzing Or22 sequences for polymorphisms that co-varied with preference for YVN over YVL and, finally, use genome editing to swap allele types (chimeric for non-chimeric) in an otherwise identical genetic background.

## Results

### Expanding the Or22 behavioral panel shows enhanced sensitivity to YVN in chimeric Or22 lines, similar to the original set

We first sought to assay the behavior of additional wild-type, fly lines to ascertain if the correlation between Or22 allele and increased sensitivity to YVN held true in a larger group. In addition to the existing panel of 14 lines, we obtained ten additional fly lines from Africa and Australia (Table 1), determined their allele type at the Or22 locus and tested their sensitivity to YVN using a trap-based olfactory assay (10). These additional lines behaved in a manner consistent with our hypothesis that the chimeric allele mediates increased sensitivity to YVN (Figure 1).

**Table 1.** ***D. melanogaster* lines used in behavioral panel**.

| *D. melanogaster* line | Collection site | Or22 genotype | Source<sup>^</sup> |
| --- | --- | --- | --- |
| CantonS | Canton, OH, USA | Non-chimeric | Eisen laboratory stock (22) |
| OreR | Roseburg, Oregon, USA | Non-chimeric | Eisen laboratory stock (22) |
| Ral437 | Raleigh, NC, USA | Chimeric | (9) |
| Ral324 | Raleigh, NC, USA | Non-chimeric | (9) |
| Ral705 | Raleigh, NC, USA | Non-chimeric | (9) |
| GRAC | Crete, Greece | Non-chimeric |  |
| GR2 | Crete, Greece | Chimeric |  |
| GR21 | Crete, Greece | Chimeric |  |
| RW1001* | Cupertino, CA, USA | Chimeric |  |
| RW1005* | Cupertino, CA, USA | Chimeric |  |
| RW1008 | Cupertino, CA, USA | Non-chimeric |  |
| RW1011 | Cupertino, CA, USA | Non-chimeric |  |
| Cellar8.3* | Healdsburg, CA | Chimeric |  |
| ME | Bowdoin, ME, USA | Chimeric | (23, 24) |
| PEN | Media, PA, USA | Chimeric | (23, 24) |
| FL | Homestead, FL, USA | Non-chimeric | (23, 24) |
| MAU9* | Rockhampton, AU | Non-chimeric | (25)<sup>#</sup> |
| MAU24* | Rockhampton, AU | Chimeric | (25)<sup>#</sup> |
| MAU31* | Rockhampton, AU | Chimeric | (25)<sup>#</sup> |
| FP6* | Sydney, AU | Non-chimeric | (25)<sup>#</sup> |
| FP8* | Sydney, AU | Non-chimeric | (25)<sup>#</sup> |
| FP16* | Sydney, AU | Chimeric | (25)<sup>#</sup> |
| CW105* | Mbengwi, Cameroon | Chimeric | (20)<sup>#</sup> |
| EZ2* | Ziway, Ethiopia | Chimeric | (20)<sup>#</sup> |
| SP90* | Phalaborwa, South Africa | Non-chimeric | (20)<sup>#</sup> |
| ZS56* | Sengwa, Zembabwe | Chimeric | (20)<sup>#</sup> |

\*Fly lines that were added to initial set examined by (21).
^blank indicates lines that were collected and established as inbred, isofemale lines for this study
<sup>#</sup>Provided by the Begun laboratory (UC Davis)

**Figure 1.**
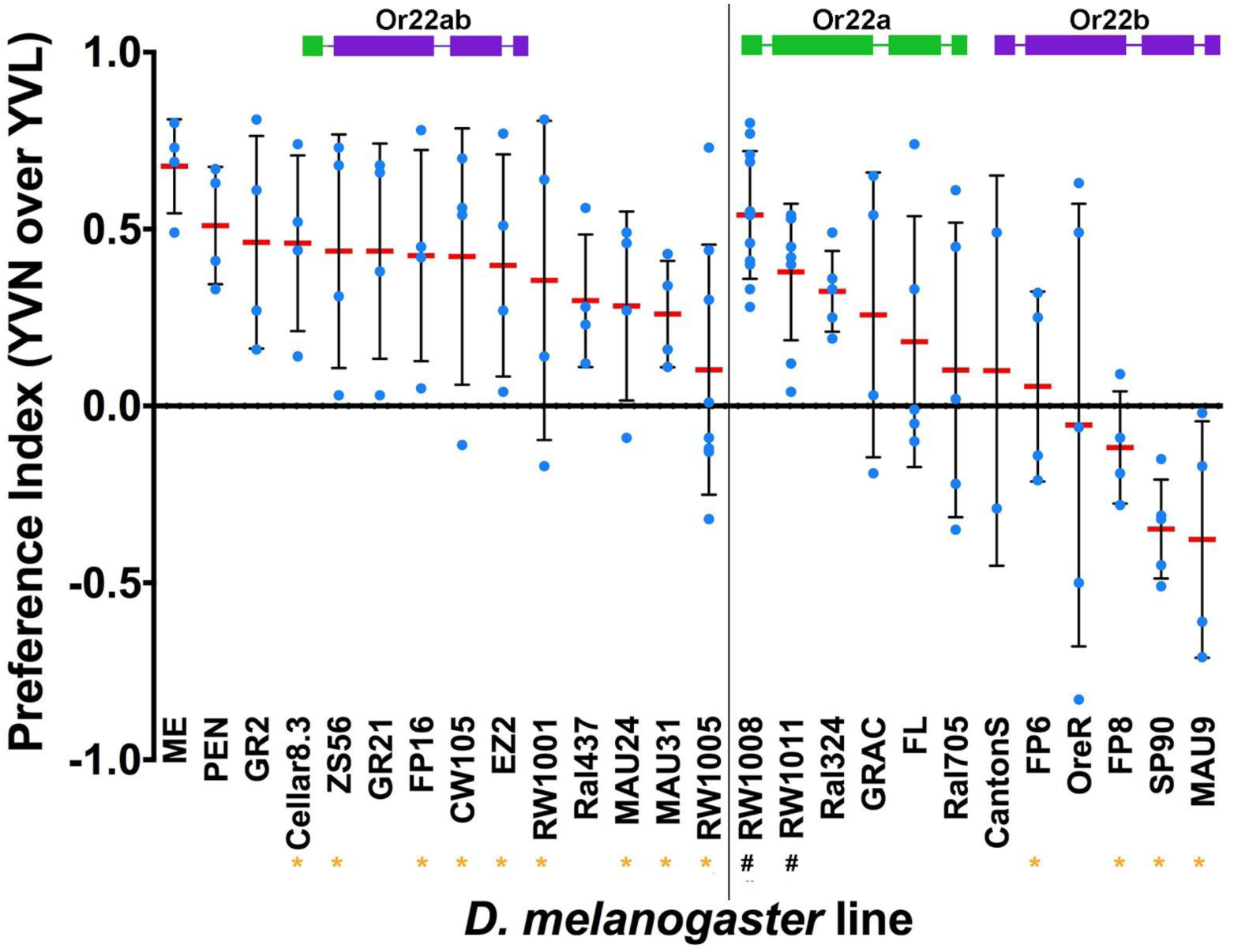
Behavior of each fly line in our Or22 behavioral panel in olfactory trap assay. Replicate behavioral experiments are plotted as blue dots; red lines indicate mean and black lines indicate standard deviation for all replicates. Positive preference index indicates a preference for yeast grown on limiting nitrogen over limiting sugar (i.e. preference for YVN); preference index of 0 indicates lack of sensitivity to YVN; negative preference index indicates preference for YVL. Lines to the left of the black vertical line have chimeric Or22 alleles (Or22ab); lines to the right have non-chimeric (Or22a and Or22b) alleles. Orange asterisks next to fly line indicate new additions to the behavioral panel. Black octothorpes next to fly line indicate that these were included in the original panel (21) but retested for this study.

### Polymorphisms in Or22 locus weakly correlate with behavioral trends

Although our expanded behavioral panel showed the same general pattern of chimeric sensitivity to YVN, there was some variability in behavior between lines with the same Or22 length variant. To determine if this variation could be attributed to nucleotide variation at the Or22 locus, we set out to clone and sequence each Or22 locus present in our behavioral panel. Although this seemed like a straightforward task, it proved to be immensely challenging. This difficulty, in fact, had been the reason why no sequence information for these alleles was available prior to this study. While the chimeric alleles can be amplified, cloned and sequenced with relative ease, non-chimeric alleles require a particular set of atypical conditions during PCR and a very large amount of template (see Methods). Additionally, Sanger sequencing across the non-chimeric alleles required different primers than for sequencing chimeric alleles, probably due to mis-priming issues in the presence of the non-chimeric tandem duplication. After much effort, we were able to clone, Sanger sequence and assemble at least three Or22 amplicons from each fly line from which we generated a line consensus that we used to called nucleotide polymorphisms (e.g. SNPs and indels). Analysis of these polymorphisms did not reveal patterns underlying sequence variants that consistently tracked with mean preference index for YVN (Figure 2).

**Figure 2.**
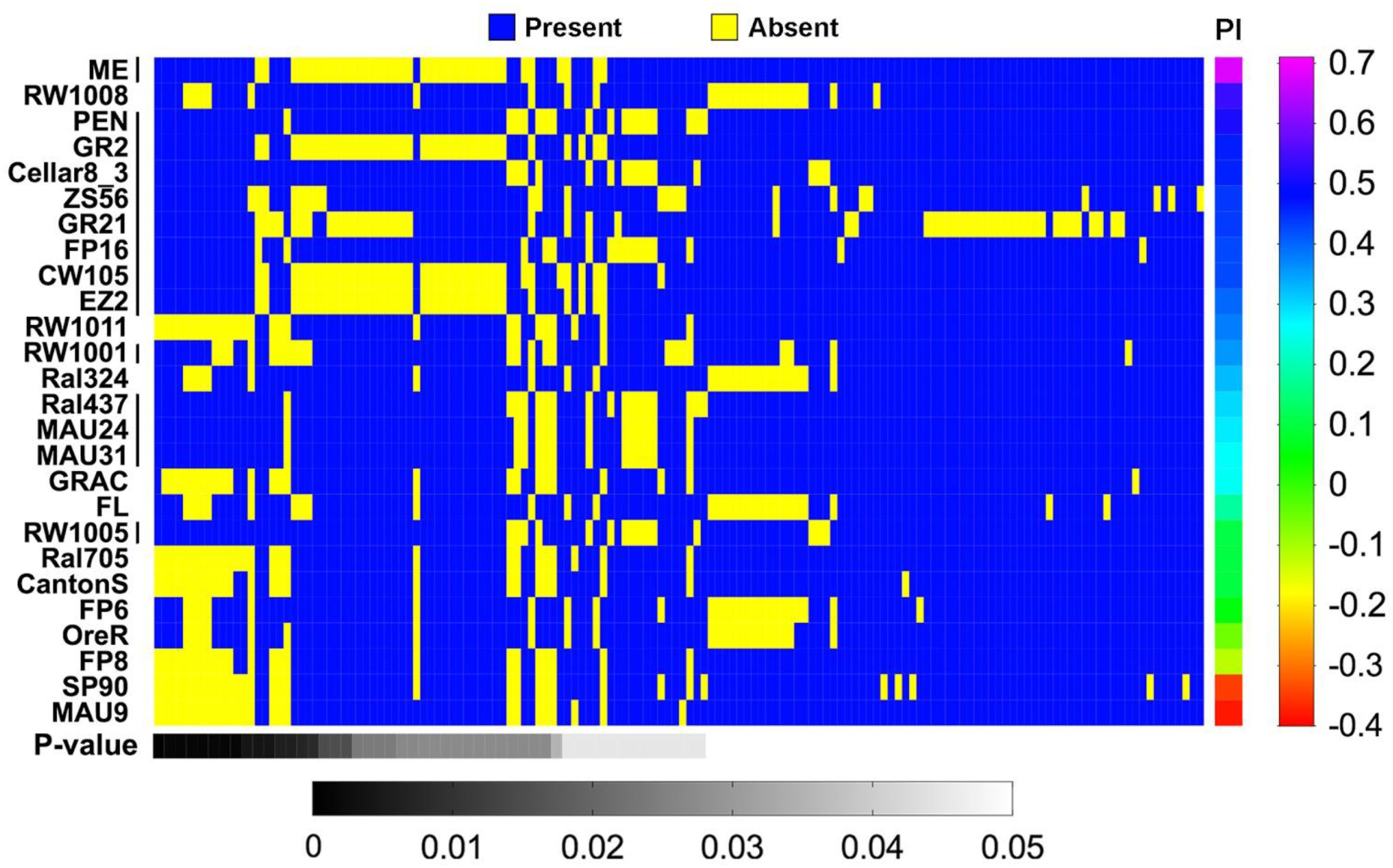
Sequence polymorphism analysis for 24 *D. melanogaster* lines in Or22 behavioral panel. For each polymorphism, a t-test was performed between the set of preference indices for YVN (PI) for lines where the variant was present or absent. Polymorphisms are ranked by p-value (shown below heatmap). No significant correlation between variance across trials for a given line and polymorphisms were found (data not shown). Fly lines are ordered by preference index (highest at top). Mean preference index for each line is given on the right. Black lines to the right of strain names indicate strains with chimeric Or22 allele. All other strains are non-chimeric.

### Replacement of a non-chimeric with a chimeric Or22 allele does not confer sensitivity to YVN

In order to directly test our hypothesis that chimeric alleles drive sensitivity to YVN, we first took advantage of the empty neuron odorant receptor system established by the Carlson group to test odorant receptor function (3). First, we cloned a chimeric allele (Ral437-Or22) into a vector under the control of UAS expression. Then, through a series of crosses, we generated flies with this UAS construct and GAL4 expression under the control of the Or22 promoter in an Or22 null background (Δhalo, (14)). Unfortunately, the Δhalo homozygotes were very sick and their health was not improved by expressing Or22 in our experimental animals. We were unable to generate sufficient numbers of animals for our behavioral assay and moved to adopt a different approach.

We next turned to the CRISPR-Cas9 gene-editing system and began implementing a two-step allelic replacement scheme (Figure S1). We opted to perform this swap in two steps rather than one due to the technical challenges we had encountered with amplification of the Or22 locus. For example, we were concerned that if we were to swap a non-chimeric allele with a chimeric allele, we would be able to robustly detect heterozygotes but would be unclear whether the non-chimeric allele was successfully removed when generating the homozygote. In the converse swapping experiment, we would have the opposite problem: detecting heterozygotes would be difficult due to the preferential amplification of the chimeric over the non-chimeric allele. In order to aid our detection of transformants, we designed the first step to replace the Or22 allele with a visible marker (beta-tubulin GFP cassette) so we could use visual screening to identify heterozygotes during the first round of replacement and homozygotes during the second.

In the first round of replacement of a non-chimeric allele (OreR) with our place holder cassette, we learned that our visible marker was not the reliable indicator of transformation that we hoped it would be. Though we expected global GFP expression in our transformed heterozygotes, we observed a weak symmetric GFP signal in the thorax and abdomen (Figure S2). This led us to identify some heterozygotes which were confirmed by non-lethal genotyping. Consistent with our expectations, the YVN sensitivity of the resultant homozygotes from these transformants phenocopied the parental line (Figure 3). We continued with our second round of replacement to swap in a chimeric allele (ME) in the place of our visible marker, obtained homozygotes (MEΔOreR) and assayed their behavior (Figure 3). Our MEΔOreR flies did not show an increased preference for YVN and so did not support our hypothesis.

**Figure 3.**
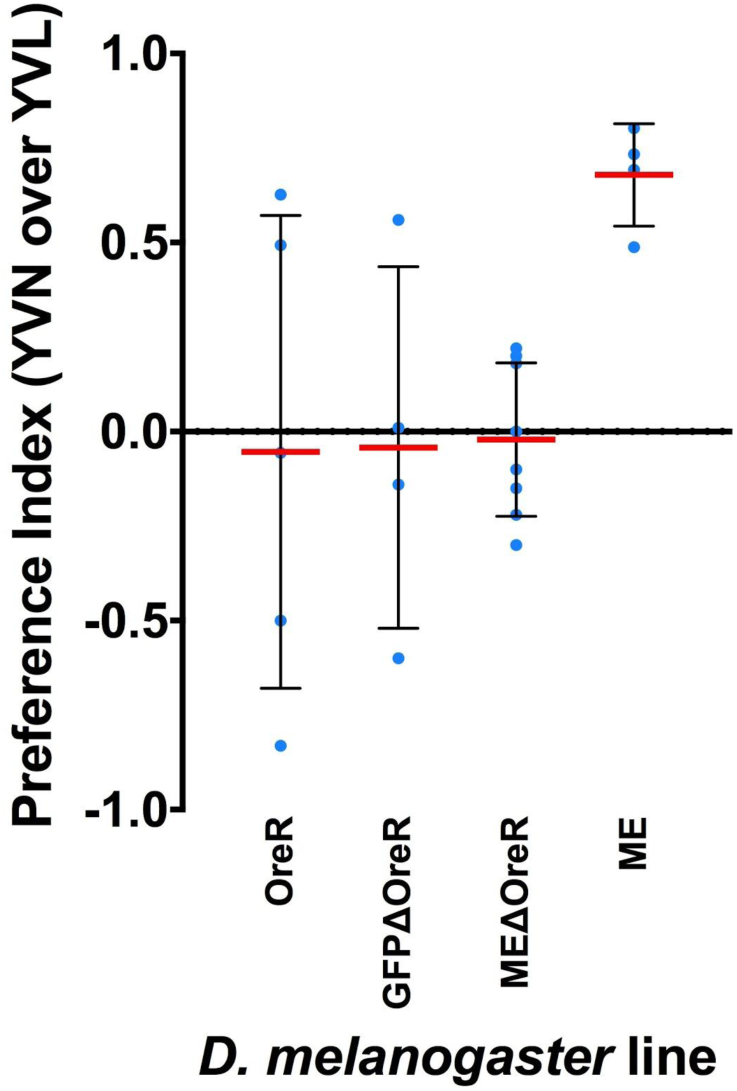
Behavior of donor lines (OreR and ME), intermediate (GFPΔOreR) and swapped line (MEΔOreR) in olfactory trap assay. Replicate behavioral experiments are plotted as blue dots; red lines indicate mean and black lines indicate standard deviation for all replicates. Positive or preference index indicates a preference for YVN; preference index of 0 indicates lack of sensitivity to YVN; negative preference index indicates preference for YVL.

### Crosses between chimeric and non-chimeric lines do not show a consistent inheritance pattern

In parallel with functional studies, we performed crosses between chimeric and non-chimeric fly lines to determine the heritability of the Or22 locus with respect to behavioral sensitivity for YVN. We first crossed two fly lines with consistent, yet strikingly different behavioral responses to YVN over YVL. The OreR fly line, homozygous for the non-chimeric allele of Or22, has no behavioral preference for yeast grown on YVN or YVL while the ME fly line, homozygous for the chimeric allele, exhibits a strong preference for YVN (Figure 1).

The OreR x ME cross was performed in both directions (i.e. one cross used an OreR virgin female and ME male; the other an OreR male and ME virgin female) and progeny were assayed for sensitivity for YVN. We found that the progeny of these crosses yielded inconsistent behavioral responses depending on the directionality of the cross (Figure 4). When ME virgin females were crossed to OreR males, the progeny preferred YVN over YVL, phenocopying the ME chimeric parental line. However, when OreR females were crossed to ME males, the F1s showed exhibited an intermediate phenotype.

**Figure 4.**
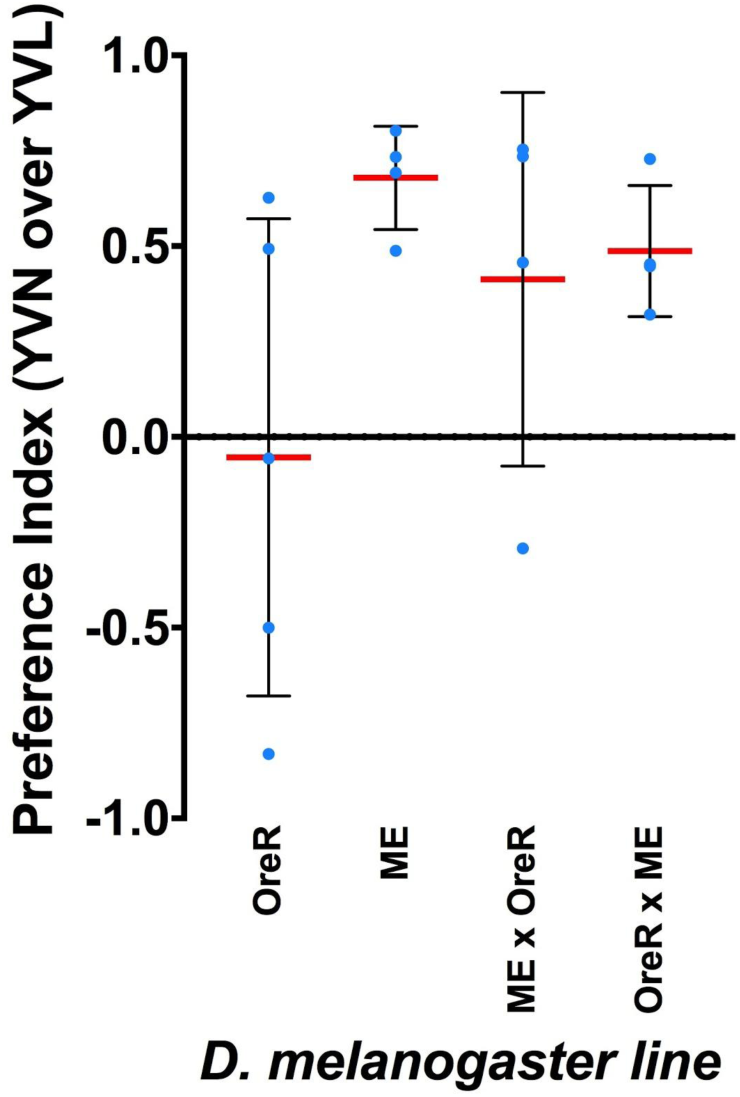
Behavioral preference for YVN over YVL for F1 crosses between ME and OreR fly lines in comparison to parental behavior. Replicate behavioral experiments are plotted as blue dots; red lines indicate mean and black lines indicate standard deviation for all replicates. Positive or preference index indicates a preference for YVN; preference index of 0 indicates lack of sensitivity to YVN; negative preference index indicates preference for YVL. Virgin females used in each cross are listed first.

The directional inconsistencies of this cross suggest that the genetics underlying the sensitivity for YVN may be sex linked. Based on our hypothesis, we did not expect a sex-linked inheritance pattern because the Or22 locus is located on chromosome 2L in *Drosophila melanogaster*. To confirm these results, we conducted additional crosses between chimeric and non-chimeric lines by crossing the chimeric Ral437 line to three different non-chimeric lines, OreR, CantonS, and Ral324. As a control, we also crossed the three non-chimeric lines to each other. The F1 progeny from each of these crosses were assayed for for sensitivity to YVN (Figure 5). Again, the observed behaviors of these flies were inconsistent with our hypothesis that the Or22 allele is responsible for mediating sensitivity to YVN.

**Figure 5.**
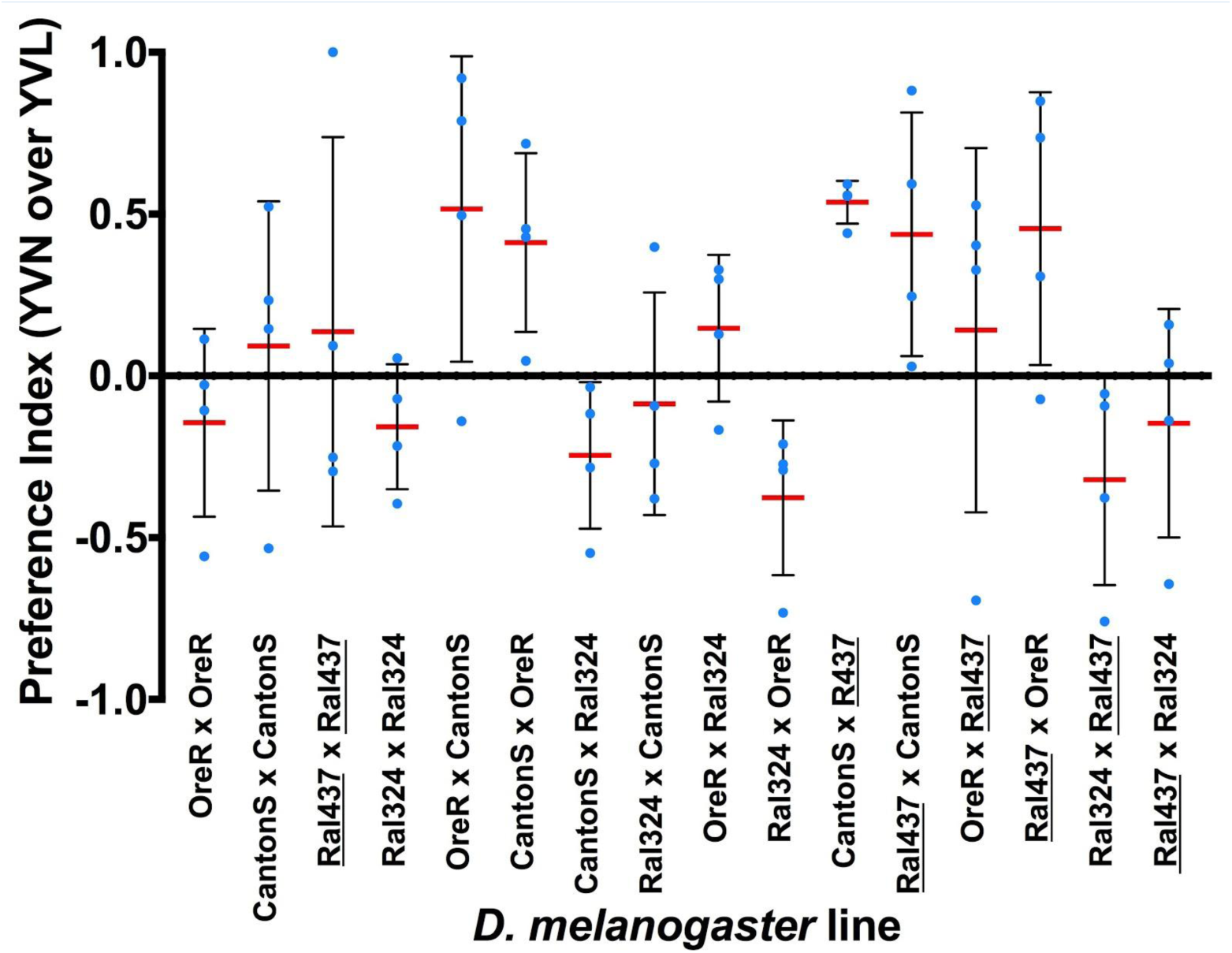
Behavioral preference for YVN over YVL for F1 crosses between one chimeric fly line and three non-chimeric fly lines. Replicate behavioral experiments are plotted as blue dots; red lines indicate mean and black lines indicate standard deviation for all replicates. Positive or preference index indicates a preference for YVN; preference index of 0 indicates lack of sensitivity to YVN; negative preference index indicates preference for YVL. Virgin females for each cross are listed first. The only chimeric line tested here is Ral437, which is denoted as chimeric by a black underline.

Within this set, we were particularly puzzled by the outcome of the Ral437 x Ral437 control cross. Previously, we had found that Ral437 preferred YVN over YVL (Figure 1), but in this experiment Ral437 x Ral437 F1s showed no preference at all. We later determined that this inconsistency was a result of our Ral437 stock having passed through a population bottleneck between the time of these assays (for various reasons, the total number of adults in our Ral437 stock plummeted after the initial assays; all progeny of the second were considerably more inbred than previously). After genotyping the Or22 locus of the stocks before and after Ral437 bottleneck, we found that the original Ral437 stock was actually heterozygous at the Or22 locus. Most flies carried the chimeric allele but the non-chimeric allele was present and maintained at low abundance. During the bottleneck, the non-chimeric allele became over-represented, thus shifting the allele frequencies of the Ral437 stock from chimeric to non-chimeric. We believe that this explains the weaker preference for YVN over YVL in the original Ral437 fly line (Figure 1) and the complete loss of preference in our subsequent cross experiment (Figure 5). Ultimately, we were unable to clarify the heritability of the Or22 locus from these data. At face value, these crosses suggest that the Or22 locus does not or is not the only locus underlying behavioral sensitivity for YVN.

## Discussion

Though we initially observed a strong correlation between increased preference to YVN and a chimeric variant of Or22 and this correlation held when expanding the number of fly lines examined, our additional follow up experiments were inconsistent with our hypothesis that chimeric Or22 alleles confer heightened sensitivity to YVN. It is notoriously difficult to link a single gene to a behavioral phenotype due to the complicated nature of behavior. Still, we hope that the data presented in this work will contribute to the efforts in the field of behavioral genetics. While it is still possible that the Or22 locus is involved in sensitivity to YVN, we offer some alternative hypotheses and additional experiments to further investigate the variation in this behavior in wild *Drosophila* lines.

### Possible epistatic interactions between polymorphisms within the Or22 locus

Though it is possible that there are epistatic interactions between polymorphisms within the Or22 locus that could significantly correlate with preference index, we postponed these analyses until we learned the outcome of the functional experiment, reasoning that if replacement of a non-chimeric Or22 allele with a chimeric one did not result in the expected behavior, these analyses would be irrelevant. As this turned out to be the case, these analyses were never performed. Still, even with the results of our functional experiment, we cannot completely discount the possibility of epistatic interactions between Or22 and another gene or genes (see below).

### Multiple loci may mediate sensitivity to YVN in *D. melanogaster*

At this juncture, it is clear that Or22 alone does not explain the variation in sensitivity to YVN in *D. melanogaster* lines. Given the sequence variation at this locus, it still seems possible that Or22 is in some way involved in attraction to yeast, though at this point we do not understand the role it plays. Given the complexity in chemical signaling between yeast and flies, it seems more likely that the molecular basis for this attraction in flies lies not in one gene but in the combined or epistatic effects of many. This hypothesis would be best addressed by taking advantage of the Raleigh line collection, a set of recently established, iso-female *D. melanogaster* lines (9). As all of these lines have been sequenced, it would be feasible to find a subset of flies that vary in their response to YVN and perform a genome-wide association study to determine sequence polymorphisms that correlate with this preference.

### Is the yeast attraction phenotype robust enough?

However, before such a study is performed, it should be considered whether the behavioral differences in the chosen panel of fly lines are consistent enough from generation to generation to make this feasible. While the number of replicate behavioral assays run was certainly appropriate for measuring previous phenotypes, (10) it may need to be increased for subsequent experiments in this line of inquiry. It is possible that different testing conditions (e.g. yeast strains) could reveal more robust behavioral differences between these lines. As is the case with many other behavioral assays, this it is certainly not the only set of conditions that could be used.

### MEΔOreR Or22 locus exhibits aberrant amplification behavior

Despite the confirmation of transformants through non-lethal PCR screening during the second round of replacement, PCR genotyping of the final homozygotes gave unexpectedly small amplification products (Figure S3). Sequencing these products and those from the screening steps prior revealed the expected sequence, with the caveat that, in the non-lethal genotyping amplicons, the sequences became heterozygous about half way through. We are hard-pressed to explain why, by all apparent measures, these animals appear to be our desired transformants and yet show this unexpected PCR phenotype. Though we believe that our transformants have the correct genotypes, we thought this was an important caveat that needs to be explored for future work on this project.

Finally, it is possible that the effects of the non-chimeric Or22 allele replacement in our functional experiment are masked by other behavioral deficiencies or phenotypes in the OreR background. OreR is a common lab fly line and its decades-long maintenance in the laboratory under unnatural conditions may have selected for behavioral phenotypes that are ecologically irrelevant or potentially conflicting with the behaviors tested here. If another Or22 allele replacement experiment such as the one described above was repeated, we suggest using a more recently established non-chimeric background line.

### Concluding thoughts

Given the importance of yeast to *D. melanogaster*, the variation in preference towards yeasts grown under different conditions that we observed in fly lines collected from around the world is likely to have ecological relevance. Understanding the basis of this variation can only improve our understanding of the complex relationship between flies and yeast and on a more general scale, how behavior is encoded in the genome. The fly behaviors underlying this relationship are likely to have multiple components, many of which can be controlled under laboratory conditions but unfortunately, never completely. These caveats are what makes studying behavior challenging and why we know so little about the genetics encoding natural behaviors. In the spirit of science, we hope that this data, although subject to the complexities of behavioral phenotypes, will still be informative and productive in generating new questions in the field.

## Acknowledgments

We’d like to thank our high school volunteers Addie Norgaard and Julie Deng for collecting behavior data for the Australian fly lines and for helping to virgin for our reciprocal crosses. We’re also grateful to the Begun lab for providing many of our fly lines, to the Carlson lab for providing flies and protocols for the empty neuron experiment and to Dr. Aguadé for sharing her Or22 amplification advice. This research was supported by MBE’s Investigator Award from the Howard Hughes Medical Institute as well as CE’s and AQ’s National Science Foundation Graduate Research Fellowships.

## Materials and Methods

### Fly stocks

All *Drosophila melanogaster* lines used in behavioral panel are shown in Table 1. Additional lines used were Attp64 (BestGene) and w; Δhalo/CyO; Or22a-GAL4/TM3 (J.R. Carlson, personal communication). All lines were reared on medium from UC Berkeley’s Koshland fly kitchen (0.68% agar, 6.68% cornmeal, 2.7% yeast, 1.6% sucrose, 0.75% sodium tartrate tetrahydrate, 5.6 mM CaCl2, 8.2% molasses, 0.09% tegosept, 0.77% ethanol, 0.46% propionic acid) supplemented with activated dry yeast pellets at 25C on a 12:12 photoperiod unless otherwise stated.

### Olfactory behavior assay

The behavior assays in this study were performed as described in (10) Briefly, *Drosophila melanogaster* lines were raised at room temperature (21-23C) on Koshland diet. Newly eclosed flies were pushed onto new food daily and aged at room temperature for at least four days under ambient lighting conditions (i.e. adjacent to a window) before being used in behavior assays.

The day prior to the start of the behavior assay, the *Saccharomyces cerevisiae* strain, I14 (26), was plated onto either YVN or YVL media (Table 2) and grown at 30C for 22 hours. Two plates for each media type were streaked out per arena.

**Table 2.**
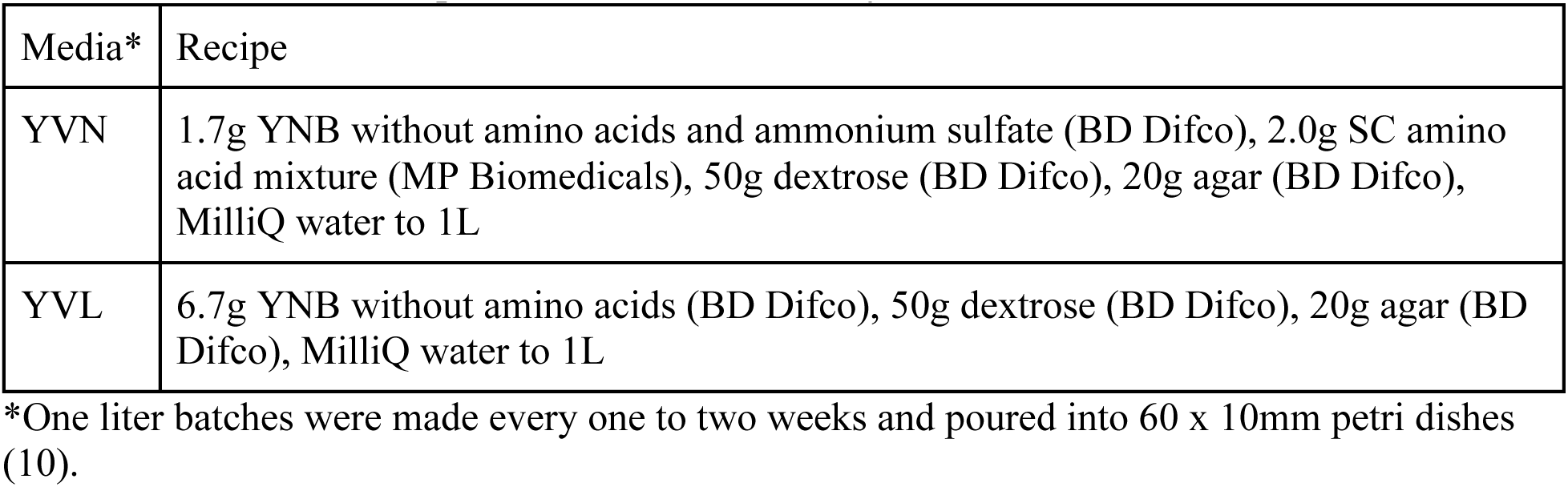
**Yeast media recipes used in behavior assays**.

| Media* | Recipe |
| --- | --- |
| YVN | 1.7g YNB without amino acids and ammonium sulfate (BD Difco), 2.0g SC amino acid mixture (MP Biomedicals), 50g dextrose (BD Difco), 20g agar (BD Difco), MilliQ water to 1L |
| YVL | 6.7g YNB without amino acids (BD Difco), 50g dextrose (BD Difco), 20g agar (BD Difco), MilliQ water to 1L |

\*One liter batches were made every one to two weeks and poured into 60 x 10mm petri dishes (10).

The following day, grown plates were removed from the incubator, fitted with a custom, 3D printed lid, and secured with Parafilm. Lids were fitted with a 50mL conical centrifuge tube (Falcon) with the end removed and covered in mesh. A funnel was fashioned from 150mm filter paper (Whatman, Grade 1) and a 5mm hole snipped off the tip. This funnel was used to top the centrifuge tube and secured with tape. Two traps for each media type were placed into behavior arenas (*Drosophila* population cages, 24” x 12” clear acrylic cylinders, TAP plastics) fitted with netting (Genesse Scientific) as shown in Figure 6. All possible orientations of YVN and YVL plates within were tested to control for environmental effects (e.g. attractivity to light).

**Figure 6.**
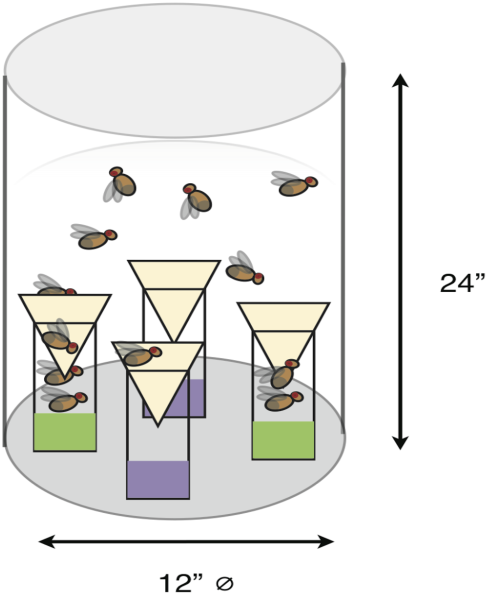
Schematic of behavior assay per (10). *Saccharomyces cerevisiae*, strain I14 (26), was grown on either YVN or YVL media and used to bait custom made traps. One hundred and twenty adult *Drosophila* (4-10 days old) of mixed sex were allowed to choose between traps over an 18 hour period. Preference was quantified by the number of flies in each trap at the end of the assay.

One hundred and twenty 4-10 day old mixtures of male and female *Drosophila melanogaster* were anesthetized with CO2 and allowed to recover on Koshland diet for 2 hours before being used in behavior assays. Flies were introduced into behavior arenas at 3pm and allowed survey traps. After 18 hours, traps were removed from the arena and the number of flies in each trap were counted, sexed and recorded. Flies were discarded after counting so flies were only used in behavior assays once. A preference index was calculated from the number of flies in each trap as follows:

> For **A** = total number of flies in YVN traps
>
> For **B** = total number of flies in YVL traps
>
> Preference Index = (**A** - **B**)/(**A** + **B**)

### Genomic DNA extraction from behavior panel fly lines

For each line, three females were pooled in a single DNA extraction using either the QIAamp DNA Micro (QIAGEN) following the manufacturer’s instructions for the isolation of genomic DNA from less than 10 mg tissue or the PureGene Tissue kit (Gentra). Concentration of each DNA sample was quantified using the Qubit High Sensitivity dsDNA kit.

### Cloning Or22 alleles via TOPO TA

Or22 alleles were cloned by amplification with GoTaq mastermix (Promega) using 240 ng of template gDNA, and 400 nM each o2F and o2R (Table 3) in a 50 uL reaction. Reactions were cycled using a specialized thermocycler protocol (M. Aguadé, personal communication): an initial melting step of 94 C for 3 min followed by 35 cycles of 96 C for 10 seconds, 55 C for 10 seconds, 65 C for 4.5 min, then a final polymerase elongation step of 65 C for 7 min. Expected bands were excised from 1% agarose gels after running at 100V and gel purified using the QIAquick Gel Extraction (QIAGEN) kit eluting in 30 uL of buffer EB. Adenosine tails were added to these fragments in anticipation of TOPO TA (Invitrogen) cloning with 5U Taq polymerase (NEB), 280 uM dNTPs in 1x standard Taq buffer (NEB) for 20 minutes at 72C. A-tailed products were then immediately cloned into TOPO TA 2.1 vector using manufacturer’s instructions. Fresh TOPO TA reactions were drop-dialyzed on 0.025 um membrane (Millipore) floated in a 100x15 mm petri dish with sterile DI water (~25 mL) for 15 minutes at RT. Drop-dialyzed TOPO TA reactions were then transformed into DH5alpha *E. coli* via electroporation, rescued immediately with room temperature SOC and outgrown 15 min at 37 C with 180 rpm shaking before plating all cells pre-warmed LB + carbencillin (100 ug/mL) agar plates. Plates were incubated overnight at 37 C. Colonies were picked and dissolved into 5 uL of LB + kanamycin (50 ug/mL) in 96-well plates. One uL of cell suspensions were then used to template 20 uL colony PCR reactions with GoTaq mastermix (Promega) using primers o2F and o2R (800 nM each) with the following thermocycler settings: initial melt at 95C for 5 min followed by 35 rounds of 95 C for 30 seconds, 51 C for 30 seconds and 72 C for 2.5 min, then a final extension step at 72 C for 10 min. Positive hits were those that gave a 2.5 kb bands when run on a 1% agarose gel. Up to five colonies for each fly line were grown overnight in LB + kanamycin (50 ug/mL) and plasmids were extracted via MiniPrep (QIAGEN).

**Table 3.**
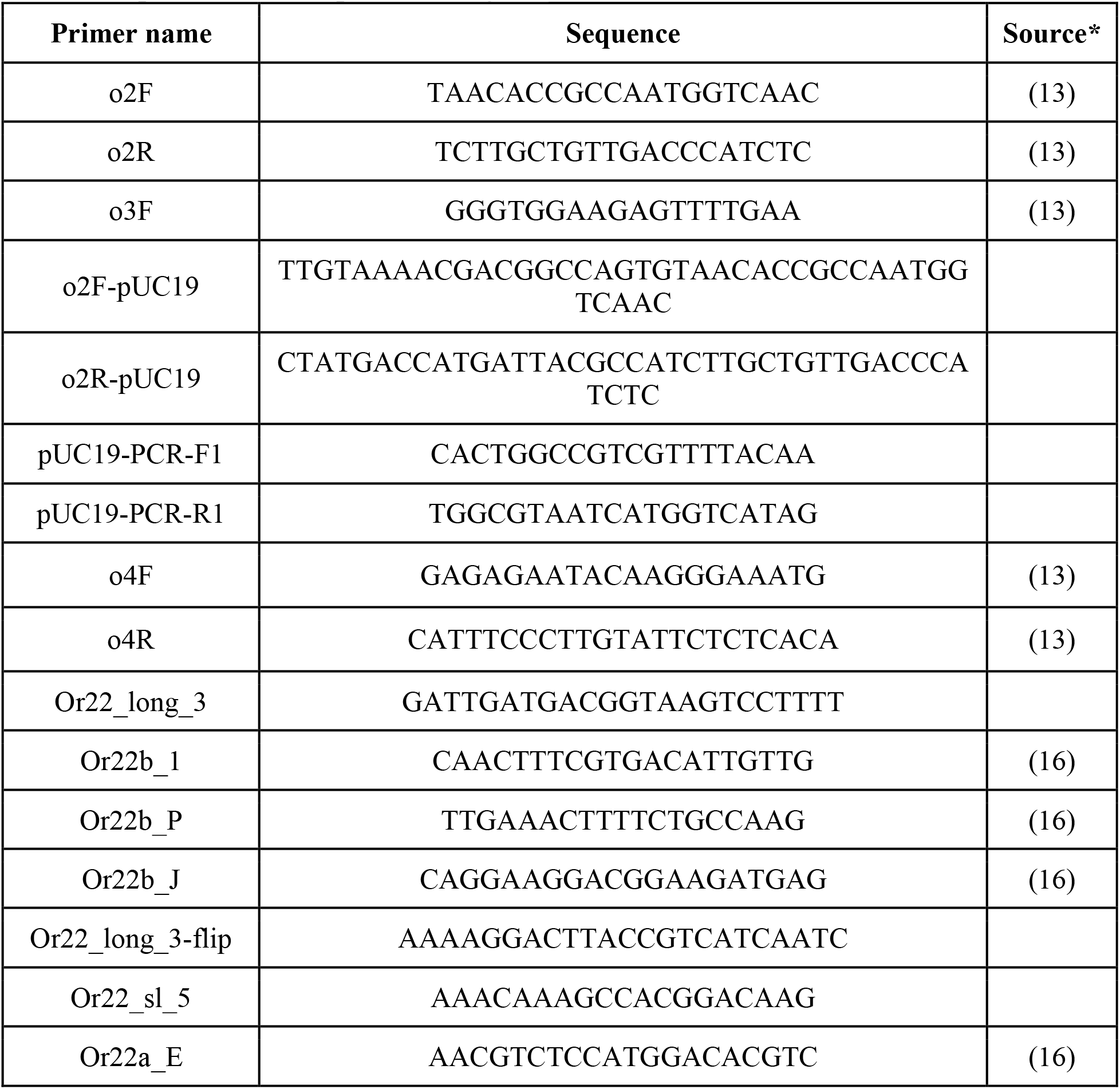
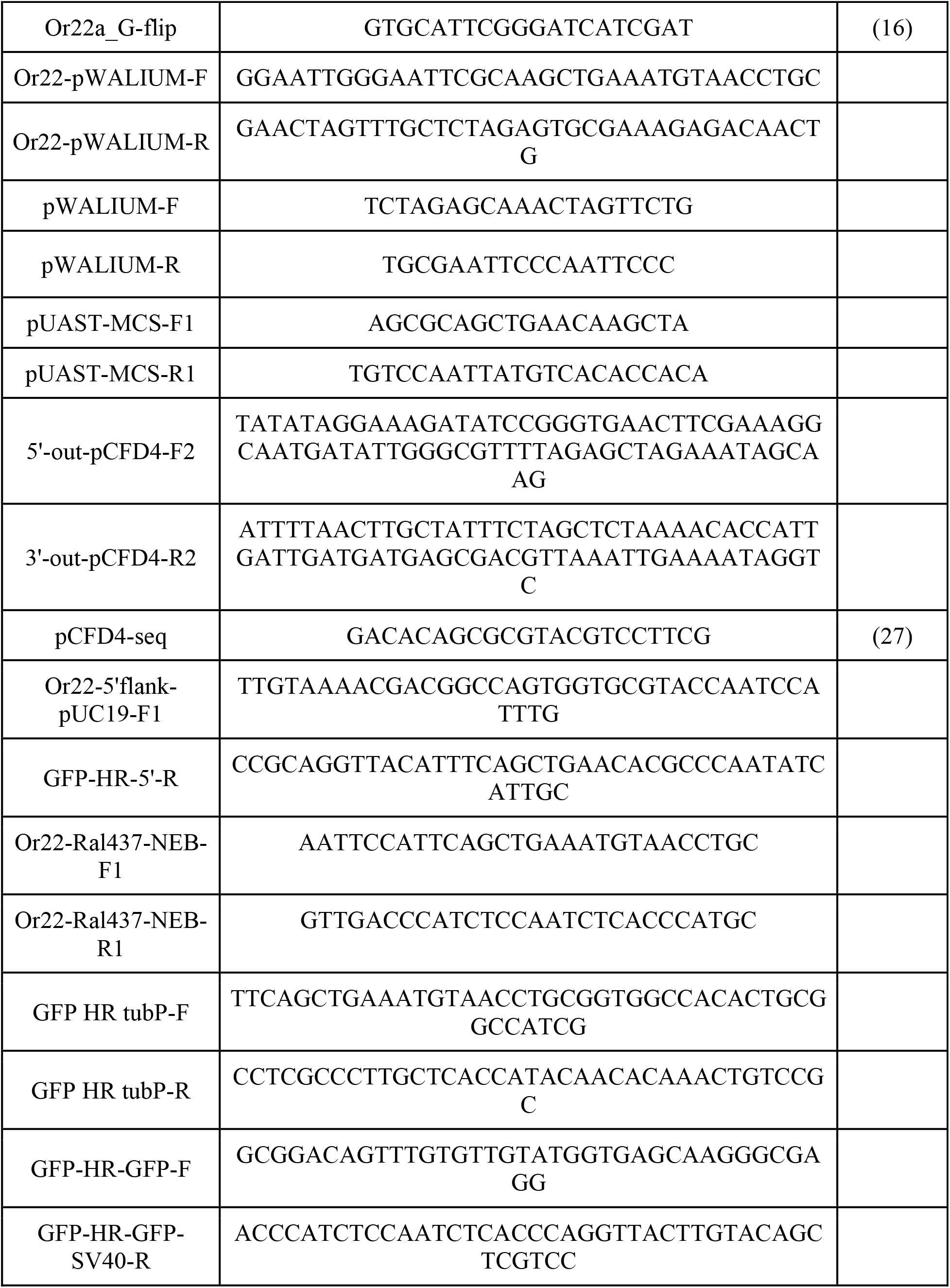
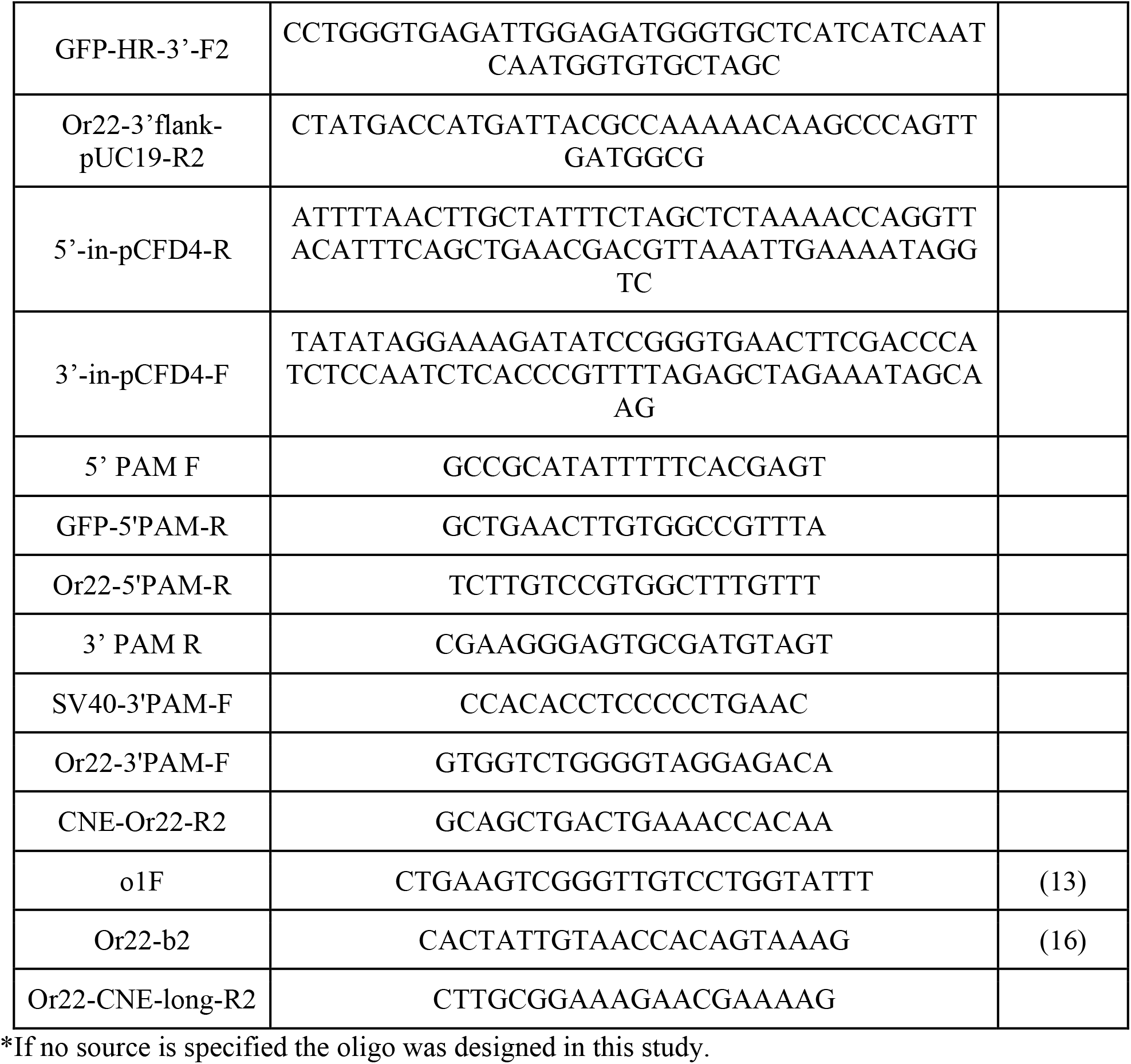
**All primers used in present study**.

### Cloning Or22 alleles via pUC19 Gibson assembly

Or22 alleles were cloned by amplification with GoTaq mastermix (Promega) using 240 ng of template gDNA, and 400 nM each o2F-pUC19 and o2R-pUC19 (Table 3) in a 50 uL reaction. Reactions were cycled using a specialized thermocycler protocol (Montserrat Aguadé, personal communication): an initial melting step of 94C for 3 min followed by 35 cycles of 96C for 10 seconds, 55C for 10 seconds, 65C for 4.5 min, then a final polymerase elongation step of 65C for 7 min. PUC19 backbone was amplified from pUC19 (Invitrogen) using pUC19-PCR-F1 and pUC19-PCR-R1 (500 nM each, Table 3) with Q5 High-Fidelity DNA Polymerase (NEB) with the following conditions: 98C for 30 sec followed by 30 rounds of 98C for 10 sec, 62C for 30 sec then 72 for 1 min, finishing with 72C for 2 min. Expected bands were excised from 1% agarose gels after running at 100V and gel purified using the QIAquick Gel Extraction (QIAGEN) kit eluting in 30 uL of buffer EB. Or22 bands were mixed with pUC19 backbone and assembled with NEBuilder HiFi DNA Assembly (NEB) incubating 1 hour at 50C but otherwise following manufacturer’s instructions. Gibson reactions were then dialyzed and transformed into DH5alpha E. coli; transformants were screened and plasmid was extracted as with TOPO TA cloning with the difference that only LB + carbencillin (100 ug/mL) agar plates were used for selection.

### Sequencing and assembling Or22 loci for each fly line in behavioral panel

Chimeric Or22 alleles were Sanger sequenced with six primers (o2F, o2R, Or22_long_3, Or22b_1, Or22b_P, Or22b_J) and non-chimeric Or22 alleles were Sanger sequenced with 10 primers (o2F, o2R, o3F, o4F, o4R, Or22_long_3-flip, Or22_sl_5, Or22b_1, Or22a_E, Or22a_G-flip) using third party services (ELIM, Barker Hall Sequencing facility) (Table 3). Loci were assembled from these sequences using SeqMan Pro (DNASTAR Lasergene v.10) after lowering signal threshold to 2 and manually checking and resolving any disagreements between reads. A consensus for each line was assembled by aligning at least three individual clones for a given fly line in SeqMan Pro.

### Polymorphism analysis for Or22 sequences

The consensus sequence for each line were aligned to the Or22 genomic reference using Geneious (version 5.1.7); chimeric sequences were split at the first intron in order to achieve alignment of the entire locus. Indels and SNPs were called manually for each consensus compared to the consensus reference of all sequenced Or22 loci to generate a presence/absence matrix of all observed polymorphisms in our set of sequenced Or22 loci. T-tests comparing the set of preference indices or variances (data not shown for latter) for all lines possessing versus lacking a given allele were performed using the stats library in Python. Data were plotted with Prism 7 (GraphPad).

### Empty neuron (Δhalo) experiment

UAS-Or22Ral437 was generated by cloning the open reading frame of Ral437 Or22 downstream of the 5x UAS in pWALIUM10 (M.R. Stadler, personal communication). To do this, Or22 was amplified from Ral437 Or22 in TOPO TA vector (Invitrogen) using primers Or22-pWALIUM-F and Or22-pWALIUM-R (500 nM each, Table 3) and pWALIUM backbone was amplified from pWALIUM using primers pWALIUM-PCR-F and pWALIUM-PCR-R (500 nM each, Table 3) with Q5 High-Fidelity DNA Polymerase (NEB) with the following conditions: 98C for 30 sec followed by 30 rounds of 98C for 10 sec, 62C for 30 sec then 72 for 45 sec (Or22) or 3:15 min (pWALIUM), finishing with 72C for 2 min. The resultant products were gel-purified, Gibson assembled (NEBuilder HiFi DNA Assembly Master Mix, NEB) transformed into chemically-competent DH5alpha *E. coli* (NEB #C2987) and selected for on LB + carbencillin (100 ug/mL) agar plates. Plasmid was extracted from 2-4 transformant clones (QIAGEN miniprep) and sequenced with pUAST-MCS-F1 and pUAST-MCS-R1 (Table 3) to confirm proper insertion had taken place. Plasmid was extracted from a verified clone (QIAGEN midiprep) and quantified using the Qubit HS dsDNA kit (Thermo Fisher Scientific). This plasmid was injected into AttP64 flies in the presence of PhiC31 recombinase and progeny were backcrossed, screened and balanced with TM3,Sb (BestGene). These w; +; UAS-Ral437Or22/TM3,Sb flies were crossed according the scheme from (3). First, UAS-Ral437Or22/TM3,Sb were crossed to w; Δhalo/Cyo; Or22a-GAL4/TM3 to generate w; Δhalo/+; UAS-Ral437Or22/TM3 and w; CyO/+; UAS-Ral437Or22/TM3 progeny. Theseprogeny were crossed to generate w; Δhalo/CyO; UAS-Ral437Or22/TM3 which were crossed back to w; Δhalo/Cyo; Or22a-GAL4/TM3 to generate w; Δhalo/Δhalo; UAS-Ral437Or22/Or22a-GAL4 flies.

### CRISPR-Cas9 Or22 allele replacement

First and second round CRISPR targets were selected using http://tools.flycrispr.molbio.wisc.edu/targetFinder/ (Table 4). Primers 5’-out-pCFD4-F2 and 3’-out-pCFD4-R2 or 5’-in-CFD-R and 3’-in-CFD-F were used to clone both sgRNAs into pCFD4 for the first or second round of CRISPR editing, respectively, per http://www.crisprflydesign.org/ (27). Constructs were verified by Sanger sequencing with pCFD4-seq (27). To generate a homologous recombination template plasmid for the first round of replacement, five fragments (pUC19 backbone (pUC19-PCR-F1 and pUC19-PCR-R1, one kilobase 5’ upstream of OreR Or22 locus (Or22-5’flank-pUC19-F1 and GFP-HR-5’-R), beta-tubulin promoter from OreR (GFP HR tubP-F and GFP HR tubP–R), GFP with SV40 3’ UTR from pGREEN-Pelican (GFP-HR-GFP-F and GFP-HR-GFP-SV40-R) and one kilobase 3’ downstream of OreR Or22 locus (GFP-HR-3’-F2 and Or22-3’flank-pUC19-R2) were amplified with Q5 High-Fidelity DNA Polymerase (NEB) using 500 nM each forward and reverse primer (Table 3) with the following conditions: 98C for 30 sec followed by 30 rounds of 98C for 10 sec, 62C for 30 sec then 72 for 2 min, finishing with 72C for 2 min. Fragments were assembled using NEBuilder HiFi DNA Assembly Master Mix (NEB), transformed into DH5alpha electrocompetent cells and plated on LB + carbencillin (100 ug/mL). Plasmid was isolated from 2-4 transformants and sequenced with primers M13F, M13R, o2F and CNE-Or22-R2 (ELIM) to confirm assembly (Table 3).

**Table 4.**
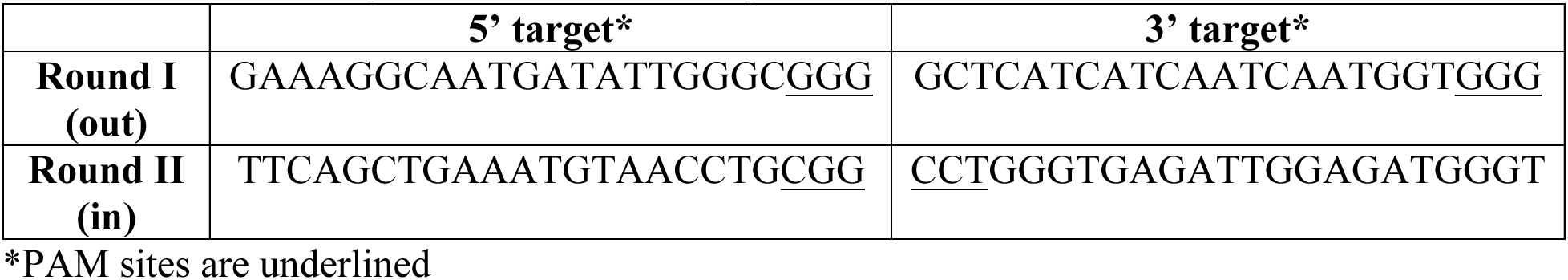
**CRISPR targets for Or22 allelic replacement**.

Plasmid from a sequence-verified clone was prepared (QIAGEN Midiprep) and quantified using the Qubit HS dsDNA kit (Thermo Fisher Scientific). OreR flies were co-injected with pHsp70-Cas9, pCFD4 containing the two synthetic guides for round I editing (outer CRISPR targets) and the GFP homologous recombination donor (Rainbow Transgenics). Injected animals were individually backcrossed to OreR; progeny were screened using a compound fluorescence microscope and by extracting DNA from pools of 50 animals from each cross (VDRC stock center protocol “Good quality *Drosophila* genomic DNA extraction”) then amplifying the 5’ and 3’ PAMs using a cocktail of primers 5’ PAM F, GFP-5’PAM-R, Or22-5’PAM-R or primers 3’ PAM R, SV40-3’PAM-F, Or22-3’PAM-F, respectively (Table 3), at a total final concentration of 1 uM for each forward and reverse primer(s) with GoTaq 2x mastermix (Promega) with the following thermocycling conditions: 95C for 5 min followed by 35 iterations of 95C for 30 seconds, 61C for 30 seconds then 72C for 30 sec then 72C for an additional 10 minutes. Sibling virgins from “hit” founder crosses were screened by non-lethal genotyping using each 5’ PAM (5’ PAM F, GFP-5’PAM-R, Or22-5’PAM-R) and 3’ PAM (3’ PAM R, SV40-3’PAM-F, Or22-3’PAM-F) primer cocktails per (28). Heterozygotes were crossed and progeny screened as above to identify homozygotes. Homozygotes were crossed, progeny were screened as above and genomic DNA from two batches of three females each was extracted with the QIAamp Micro kit (QIAGEN) then PCR genotyped and Sanger sequenced using three sets of primers to confirm homogeneity: 5’ PAM (5’ PAM F, GFP-5’PAM-R, Or22-5’PAM-R), 3’ PAM (3’ PAM R, SV40-3’PAM-F, Or22-3’PAM-F) and whole locus (5’ PAM F, 5’PAM-R) PAM primer sets and 5’PAM/3’PAM. Sibling flies were propagated as GFPΔOreR.

An analogous process was used for the second round of editing; this time pCFD4 contained the synthetic guides for round II (inner) CRISPR targets and the donor plasmid contained the Or22 allele from ME sandwiched between 5’ and 3’ Or22 flanking regions (assembled identically to the first round homologous donor template instead using four fragments (pUC19 backbone (pUC19-PCR-F1 and pUC19-PCR-R1, one kilobase 5’ upstream of OreR Or22 locus (Or22-5’flank-pUC19-F1 and GFP-HR-5’-R), ME Or22 locus (Or22-Ral437-NEB-F1 and Or22-Ral437-NEB-R1) and one kilobase 3’ downstream of OreR Or22 locus (GFP-HR-3’-F2 and Or22-3’flank-pUC19-R2) (Table 3)).These constructs and pHsp70-Cas9 were co-injected into GFPΔOreR (Rainbow Transgenics). Injected animals were individually back-crossed to GFPΔOreR then screened and homozygosed as above to establish line MEΔOreR.

### Chimeric and non-chimeric crosses

Fly lines were raised at 25C and virgins and males were collected twice a day. After five days, virgins were confirmed. Five females of a single fly line were crossed to five males of another line. Three replicates of each cross were set up and crosses were performed in both directions. As a control, virgins and males of parental fly lines were collected and crossed in parallel. F1 progeny were collected and aged for behavior assay.

